# *marApp*: An R package and web portal to calculate mutations- and genetic diversity-area relationship for conservation

**DOI:** 10.1101/2025.09.09.675155

**Authors:** Meixi Lin, Kristy Mualim, Oliver Selmoni, Moises Exposito-Alonso

## Abstract

International conservation policies including the Kunming-Montreal Global Biodiversity Framework now consider genetic diversity of wild species in their targets. However, scalable, theory-driven tools to assess and predict genetic diversity loss are still emerging, limiting their use in conservation planning. Analogous to the species-area relationship, recent work has shown that genetic diversity scales with area, described by the mutations–area relationship (MAR) or genetic–diversity–area relationship (GDAR), can approximate the genetic diversity loss in a species from habitat reduction. To enable application of these insights for conservation practitioners and policy makers, we present the *mar* R package and *marApp* web portal, a fast and user-friendly tool for MAR/GDAR analysis. The *mar* package connects genetic diversity patterns in space to ecological theory, and automates steps from reading genetic and geographic data to simulating various habitat extinction scenarios using MAR/GDAR. The *mar* package estimates only short-term genetic diversity, providing a lower-limit for the far greater long-term genetic losses, underscoring the need to maximize present-day habitat conservation before it becomes irreversible. As a case example, we showcase predictions of coral (*Acropora* sp.) genetic diversity loss from reef coverage impacts. We demonstrate the usage of *marApp* without requirements for prior genetics or coding experiences, and provide downloadable reports for conservation.

## Introduction

Genetic diversity, the amount of genetic variations within species, is the most fundamental element of biodiversity (May 1994; Gaston 2010). Genetic diversity provides the raw material on which evolutionary forces act, and is key to species adaptation to new environments and survival in small populations (Frankham 1995). However, genetic diversity has long been overlooked in international conservation policies (Laikre et al. 2020). Only in 2022, the Kunming – Montreal Global Biodiversity Framework (GBF) adopted the target “to maintain and restore the genetic diversity within and between populations of native, wild and domesticated species to maintain their adaptive potential” (CBD 2022). While genetic data is now available for thousands of species and millions of individuals, a significant gap remains within many biodiversity hotspots (Paz-Vinas et al. 2023), highlighting the need for theories and scalable approaches to quantify existing genetic diversity and predict diversity loss across species at landscape scales with simple and interpretable percentage-based estimates. However, these approaches are still in their nascent phases (Leigh et al. 2021; Hoban et al. 2023), limiting the capacity for evidence-based conservation decision-making.

In conservation practices, the macro-ecological framework of species-area relationship (SAR) has been widely applied to estimate rates of species diversity loss (Matthews and Whittaker 2015). Interestingly, this principle of “commonness of rarity” applies to not only species in a community but also mutations in a population (Hu, He, and Hubbell 2006; Exposito-Alonso et al. 2022). This fundamental similarity between theoretical population genetics and neutral macroecology led to the discovery of scaling relationships between genetic diversity and area, providing theoretical foundations for genetic diversity estimates and predictions in conservation (Exposito-Alonso et al. 2022; Mualim et al. 2024). Known as the mutations-area relationship (MAR) or the genetic-diversity-area relationship (GDAR), genetic diversity, either measured as the number of polymorphic/segregating sites (M), or the nucleotide diversity (θ_π_ or GD), increases with the area surveyed (A), following a power-law function: *M = cA*^*z*^ and *θ*_*π*_ *= cA*^*z*^. The power of this simple mathematical approach, is that upon rapid habitat area loss, one can predict the proportional reduction of genetic diversity by rearranging the MAR equation: *1-(1-A*_*loss*_)^*z*^ (Exposito-Alonso et al. 2022).

We introduce the main modules in the *mar* package using the global *Arabidopsis thaliana* example: from importing data, fitting species abundance distributions for genetic diversity, to calculating the scaling relationship between genetic diversity and sampled area, and simulating genetic extinction upon habitat loss. To enhance accessibility, we developed *marApp*, a shiny-based web portal that enables users to explore MAR/GDAR calculations and run extinction simulations on their own datasets, with step-by-step explanations. It also generates downloadable conservation reports to help managers estimate genetic diversity loss or set habitat protection targets with a single click.

### The *mar* package

The R package *mar*, available on GitHub (https://github.com/meixilin/mar), implements methods for calculating mutations-area relationships (MAR) and genetic-diversity-area relationships (GDAR). Additionally, tools to fit species-abundance distributions (SAD) in genetic data are also provided. The *mar* package depends on the *SeqArray* package (≥ v.1.28.0) for genomic data input and manipulation (Zheng et al. 2017), the *raster* package for spatial analysis and sampling (Hijmans 2025), the *sars* package for fitting MAR/GDAR relationships (Matthews et al. 2019), and the *sads* package for SAD model fitting (Prado, Dantas Miranda, and Chalom 2024).

Documentation and examples can be found on GitHub.

The workflow in *mar* is implemented through the `MARPIPELINE` function, which comprises six main steps (**Fig. 1**): (1) `data`: read genotype and coordinate files, (2) `gm`: optional data subsetting, and create `genomaps` data structure to be used for the `sfs`, `mar` and `ext` steps, (3) `sfs`: calculate the site-frequency spectrums (SFS) and fit species abundance distribution models on genetic data, (4) `mar`: build the MAR/GDAR relationship by sampling the genomaps object, (5) `ext`: simulate the extinction process on the genomaps object and predict genetic diversity loss, and (6) `plot`: automatic plotting of `MARPIPELINE` steps. Apart from the `plot` step, which requires analysis steps executed beforehand, each step in the `MARPIPELINE` is fully modularized and can be executed separately for fine-tuning and time-saving. Below we introduce the main steps and core functions in `MARPIPELINE`.

**Fig. 1.**
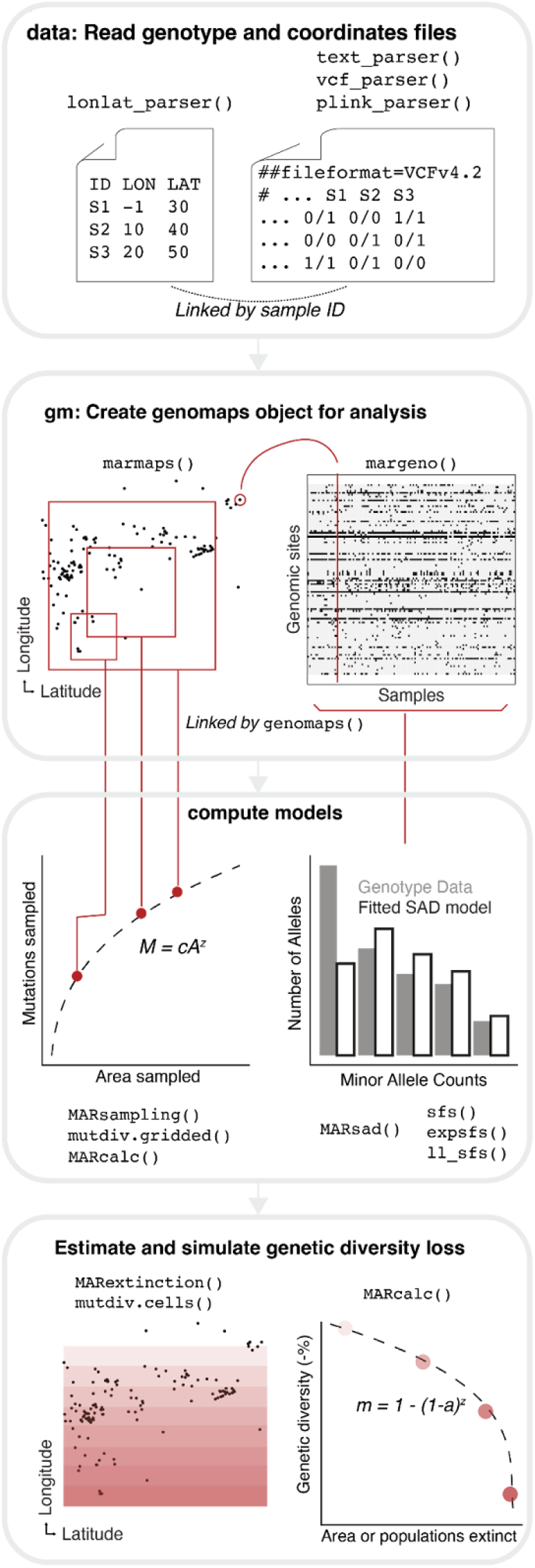
Overview of the R package *mar*. The main steps in the workflow function `MARPIPELINE` are shown. All functions provided by the *mar* package are grouped by the workflow steps.

### Data processing and genomaps object creation

The `data` step in `MARPIPELINE` requires two essential data components: coordinate and genotype of the sequenced samples. The coordinate file should be a 3-column text file containing sample identifier, longitude, and latitude (processed via `lonlat_parser` function). Genotype data can be provided in three formats: plain text files formatted as a matrix of alternative allele counts by sample and genomic position (`text_parser`), VCF files (`vcf_parser`), or PLINK files (`plink_parser`). To avoid model assumption violation, we require that genotypes should only consist of segregating bi-allelic single nucleotide polymorphisms (SNP) without missing data. Although non-diploid datasets are allowed, we recommend caution in the interpretations of MAR/GDAR results in non-diploid organisms.

In the `gm` step of `MARPIPELINE`, optional variant and sample filtering based on variant or sample identifiers can be performed. The `marmap` function then processes the filtered coordinate data to generate a raster map, with each cell value representing the sample count per location. Map resolution and coordinate reference system (CRS) can be specified manually or determined automatically using equation 12 in (Hengl 2006) and the WGS84 CRS. The `margeno` function organizes the filtered genotype data, using an S3 data structure inspired by the `SeqVarGDSClass` in *SeqArray*. Both `marmaps` and `margeno` functions create their respective S3 objects with dedicated plot and print methods. Finally, the `genomaps` function links the `marmaps` and `margeno` objects, while verifying sample identifier consistency, creating the core data structure for downstream analyses.

### Site-frequency spectrum computation

The `sfs` step of `MARPIPELINE` provides a basic summary of genetic data. The `sfs` function summarizes the genotypes data in the S3 `margeno` object to a site frequency spectrum, which is a histogram of mutation abundances, and returns an S3 `sfs` object. We also implemented the expected SFS under population genetics neutral coalescence (noted as the neutral SFS below) through the `expsfs` function, which also returns an S3 `sfs` object.

The `MARsad` function in the MAR package provides full functionality to fit popular species-abundance distribution (SAD) models, including the broken stick, geometric, lognormal, log-series, neutral metacommunity, and weibull distributions to the genotypes data through the `fitsad` function in the *sads* package (Prado, Dantas Miranda, and Chalom 2024). For SAD model predictions, to align with the more commonly used SFS visualization in population genetics, we used an in-house function (`.sadpred`) to generate the expected SFS given the inferred SAD parameters. The `MARsad` function returns a S3 `marsad` object, consisting of the inferred SAD models, the Akaike Information Criterion (AIC) table of SAD models, and the expected SFS. To facilitate comparisons of SAD-derived SFS, neutral SFS and the genotypes data SFS, we implemented a Poisson likelihood function `ll_sfs` from the ∂a∂i python package (eq. 4 in (Gutenkunst et al. 2009)).

### Geo-sampling of genetics data and building the MAR/GDAR

Building upon the observed summaries of genetic data, we proceed to empirically sample the geographical range of the data, and construct the mutations/genetic diversity-area relationships. In the `mar` step of `MARPIPELINE`, the `MARsampling` function samples genetic diversity across varying spatial scales and returns an S3 `marsamp` data frame. Based on the raster sample map (`marmaps` object) generated during the `gm` step, `MARsampling` creates replicated square subsamples of the population distribution, ranging from a single raster cell to the full range size. To account for potential geographic subsampling biases, we implemented five distinct sampling schemes, including random, directional (south-to-north and north-to-south), and focal point-based (inward and outward) methods, using a probability sampling process based on the beta distribution. Area calculations employ two strategies: *Asq* (square area in degree^2^) and *A* (cell area in km^2^). *Asq* represents the total square area defined by minimum and maximum latitude/longitude values, while *A* calculates the sum of occupied grid cell areas using the `area` function from the *raster* package. For genetic diversity, `mutdiv.gridded` implements four metrics: number of segregating/polymorphic sites (*M*), endemic segregating sites (*E*), Watterson’s θ (*θ*_*w*_), and nucleotide diversity (*θ*_*π*_). Finally, the `MARcalc` function fits the power-law model *M = cA*^*z*^ using the `sar_power` function from the *sars* package to establish the MAR/GDAR relationship for selected diversity and area metrics.

### Extinction simulation and genetic diversity loss prediction

The MAR framework can be applied in reverse to project immediate genetic diversity loss associated with rapid habitat reduction. This relationship follows the power-law equation *M = cA*^*z*^. If we note the proportion of area lost as *a*, and the proportion of genetic diversity lost as *m*, the MAR can be rearranged to predict the remaining proportion *m* of genetic diversity: *M(1-m) = c[A(1-a)]*^*z*^, which yields the genetic diversity loss prediction: *1 – m = (1 – a)*^*z*^.

In the extinction step of `MARPIPELINE`, the `MARextinction` function simulates the loss of genetic diversity, and returns an S3 `marextinct` data frame. Based on the raster sample map generated during the `gm` step, `MARextinction` removes raster cells iteratively, modeling local extinction scenarios where habitat becomes unavailable, and tracks genetic diversity in the remaining samples. Similar to the `MARsampling` function, `MARextinction` allows five extinction schemes (random, south-to-north, north-to-south, inwards and outwards), and computes four genetic diversity metrics. However, unlike `MARsampling`, individuals are sampled directly from occupied grid cells because square grids can no longer be constructed during the iterative removal. Therefore, results based on *Asq* and *A* in `MARextinction` are similar (Fig. S3). Finally, the `MARcalc` function then fits the power-law model to establish MAR/GDAR-based predictions for the proportion of remaining genetic diversity and area loss (**Fig. 1**).

### Working example: global *Arabidopsis* 1001G dataset

To showcase the *mar* package usage through command-line, we analyzed the mutations-area relationship using the accompanying data example provided in the *mar* package: `gm1001g`. This example dataset is derived from the Arabidopsis 1001 Genomes Project (1001 Genomes Consortium 2016). We downloaded the imputed SNP matrices, and accession metadata from https://1001genomes.org/ on 12-23-2024. A total of 1004 geo-referenced samples from the native continental range (excluding the USA, Canary Islands or Japan samples) were kept, and 10,000 biallelic polymorphic SNPs were randomly sampled from chromosome 1.

The analysis can be executed using a single command:

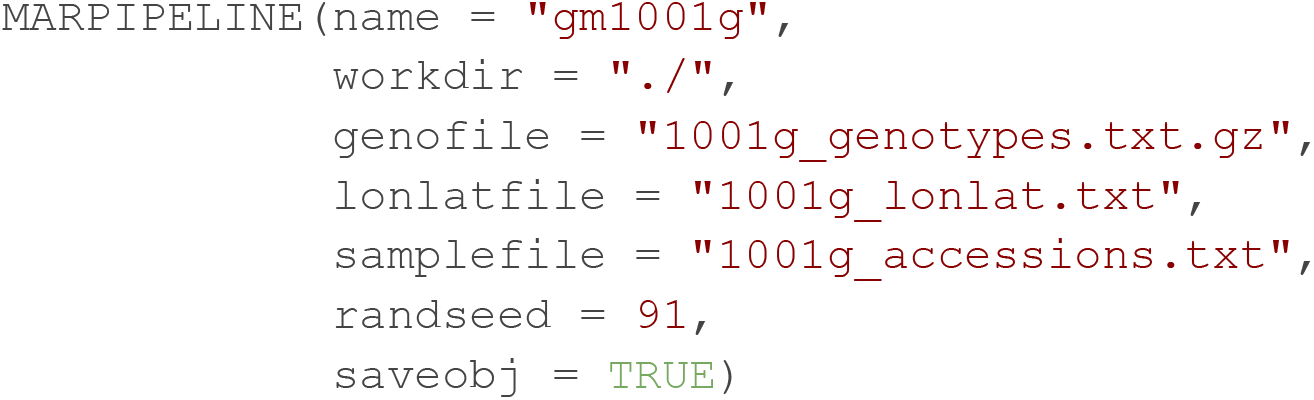

The random seed does not have to be set, we set it as 91 here for reproducibility in the results described below. `MARPIPELINE` command outputs key R Data files and PDF-formatted figures while displaying all primary results in the log file (**Table 1**).

**Table 1.** MARPIPELINE output specifications.

| Filename | Data class | Explanation | marstep |
| --- | --- | --- | --- |
| genodata_<name>.rda | S3 margeno | Genotype data imported from text files | data |
|  | character | Path to GDS file imported from VCF or PLINK files | data |
| mapsdata_<name>.rda | data.frame | Coordinates data imported from text file | data |
| gm_<name>.rda | S3 genomaps | Filtered and aligned genotype-coordinate data for analysis | gm |
| genosfs_<name>.rda | S3 sfs | Site frequency spectrum constructed from S3 margeno | sfs |
| marsad_<name>.rda | S3 marsad | MARSad output: SAD models, AIC values, and expected SFS | sfs |
| sadsfs_<name>.rda | list | All SFS outputs: data SFS, neutral SFS and SAD-predicted SFS | sfs |
| mardflist_<name>.rda | list of S3 marsamp | MARSampling outputs organized by specified sampling scheme | mar |
| marlist_<name>.rda | list of S3 marcalc | MARcalc outputs from S3 marsamp, sorted by scheme | mar |
| extdflist_<name>.rda | list of S3 marextinct | MARExtinction outputs organized by specified extinction scheme | ext |
| extlist_<name>.rda | list of S3 marcalc | MARcalc outputs from S3 marextinct, sorted by scheme | ext |
| gm_<name>.pdf | N/A | Plot of the raster sample map stored in S3 marmaps | plot |
| sfs_<name>.pdf | N/A | Plot of all SFS outputs: data SFS, neutral SFS, SAD-predicted SFS | plot |
| mar.<sch>_<name>.pdf | N/A | Plot of MAR generated from S3 marsamp with given <sch> scheme | plot |
| ext.<sch>_<name>.pdf | N/A | Plot of extinction forecasts generated from S3 marextinct | plot |

Examining the results, we found that the pipeline successfully creates a `genomaps` object, with an automatically determined spatial resolution of 0.9 degree (**Fig. 2A**). A total of 264 raster cells were created for 1004 geo-referenced *Arabidopsis* samples, with a median of two samples and a maximum of 75 samples falling in the same raster cell. In the sfs step, the SFS constructed from genotype data shows a pattern of commonness of rarity, with most SNPs segregating at lower frequencies (**Fig. 2B**). Comparing all SAD models, the Fisher’s log-series distribution (AIC = 91442.1) and the metacommunity zero-sum multinomial distribution (AIC = 91442.4) fit the data best, while the MacArthur’s broken stick distribution (AIC = 114744.1) performed the worse (**Fig. S1**). In the mar step, the mutations-area relationship was successfully constructed from spatial sampling (**Fig. 2C**). Using the random spatial sampling method, the scaling factor *z* for the number of segregating sites and area is 0.50 when using occupied grid cell areas (*A, p <* 0.001, *R*^*2*^ *=* 0.97), and 0.39 when using the total square area (*Asq, p <* 0.001, *R*^*2*^ *=* 0.46; **Table S1**). In the extinction step, habitat loss simulations were performed (**Fig. 2D**). Using the random habitat extinction method and occupied grid cell areas (*A*), the remaining genetic diversity is expected to decline as habitat becomes unsuitable, following the expression: *1-m = (1 – a)*^*0*.*38*^ (*p <* 0.001, *R*^*2*^ *=* 0.98). Note here the scaling factor *z =* 0.38 is constructed from the extinction simulation. While similar to the scaling factor constructed during the MAR sampling step (*z* = 0.50 using *A* and *z* = 0.39 using *Asq*), it is not the same value. For example, if we assume *a =* 38% global terrestrial area has been transformed according to the Millennium Ecosystem Assessment (MEA 2001) and becomes unsuitable for *Arabidopsis*, then approximately 83% of genetic diversity, measured in the number of segregating sites, still remains in the population. Once more we note that this prediction estimates current (short-term) loss, while genetic drift will continue eroding genetic diversity over time.

**Fig 2.**
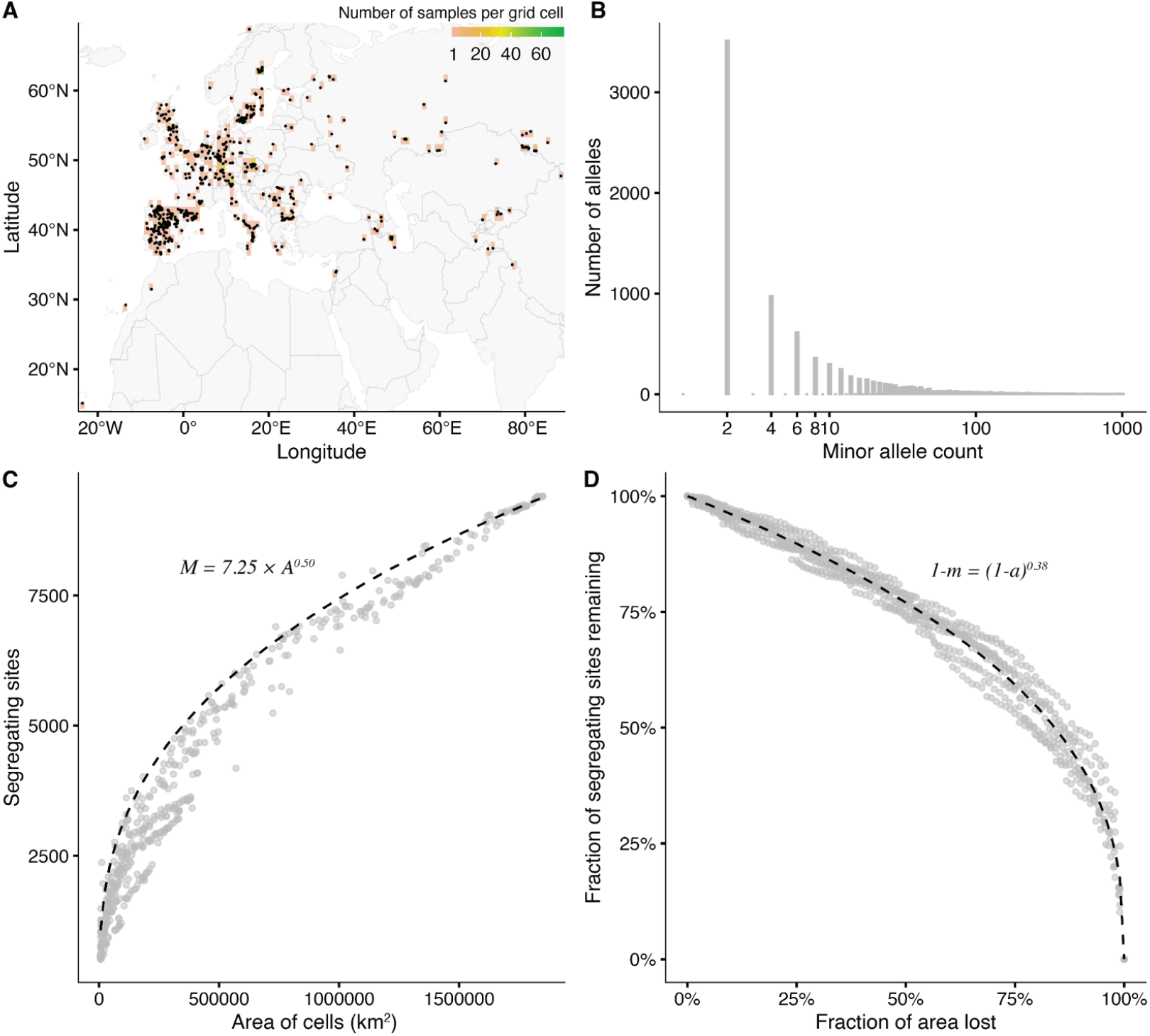
MARPIPELINE outputs from the *Arabidopsis thaliana* example. (A) Raster sample map comprising 1004 geo-referenced samples, with points indicating sample locations and color intensity reflecting sample density per cell. (B) Site frequency spectrum (SFS) of the data. (C) MAR derived from `MARsampling`, displaying segregating sites (*M*) against cell area (km^2^, *A*) with fitted *M = cA*^*z*^ curves. (D) Genetic diversity loss predictions from `MARextinction`, with fitted *1-m = (1 - a)*^*z*^ curves. Panels (B-D) are truncated, for complete results, see **Figs. S1-S3**.

Finally, we examine additional genetic diversity metrics. Akin to the species-area relationship (SAR), the mutations-area relationship was derived to approximate a power law from process-based approaches. However, following SAR, researchers have applied power law diversity-area concept to other species diversity metrics, e.g. Simpson’s diversity or other taxonomic levels (Ma 2018). We have also explored and implemented several genetic diversity metrics, including the endemic segregating sites (*E*), Watterson’s θ (*θ*_*w*_), and nucleotide diversity (*θ*_*π*_), to establish the genetic-diversity-area relationship (GDAR). A phenomenological scaling relationship between such genetic diversity metrics and area still holds, but the scaling factor *z* and *R*^*2*^ changes (**Table S1, Figs. S2, S3**). Endemic segregating mutations, which are mutations that only segregate in the sampled regions, saturates slower with area than segregating mutations (*z* = 1.28 with *A*, 0.81 with *Asq*). Endemic mutations carry characteristic information for local genetic diversity but converge to allelic richness (*M*) as the study range expands to a species’ full range. Watterson’s θ quantifies the number of segregating sites with a sample size scaling factor, yielding a faster saturation rate of genetic diversity (*z* = 0.25 with *A*, 0.15 with *Asq*) compared to *M*, the number of segregating sites. Lastly, nucleotide diversity quantifies the average pairwise differences between genetic samples. Being less sensitive to rare alleles, *θ*_*π*_ saturates the fastest with area (*z* = 0.10 with *A*, 0.07 with *Asq*), and its relationship is noisier although significant (*R*^*2*^ = 0.50 with *A*, 0.22 with *Asq, p <* 0.001).

### The *marApp* and Florida coral reef conservation example

To facilitate the use of MAR/GDAR in conservation settings, we developed a shiny-based application that implements the `MARPIPELINE` function described above, available at https://moiexpositoalonsolab.shinyapps.io/marApp/. To facilitate result interpretation, and provide background information, an introductory snippet is included in the *marApp*. We showcase visualizations with an example dataset derived from a global *Acropora* genetic meta-analysis (Selmoni, Cleves, and Expósito-Alonso 2024). This dataset includes 76 *Acropora cervicornis* (staghorn coral) individuals sampled in the Florida Reef Tract, and sequenced with Genotyping by Sequencing technology (Drury and Lirman 2021; Drury et al. 2019).

In the “Upload data” tab, select “Custom” dataset, we uploaded a text file containing *Acropora* sample index, longitude and latitude information in the “Coordinate file” section and a text file containing genotypes in the “Genotype file” section following the instructions listed in *marApp*. After both files are successfully uploaded, click the “Load data” button. A preview for the uploaded coordinate and genotype data occurred, summarizing the data input. Additionally, an interactive map created by *leaflet* was available to examine the sample locations and marmaps object created. In this example, samples mapped along the Florida Reef Tract, suggesting successful `genomaps` object creation (**Fig. 3**).

**Fig 3.**
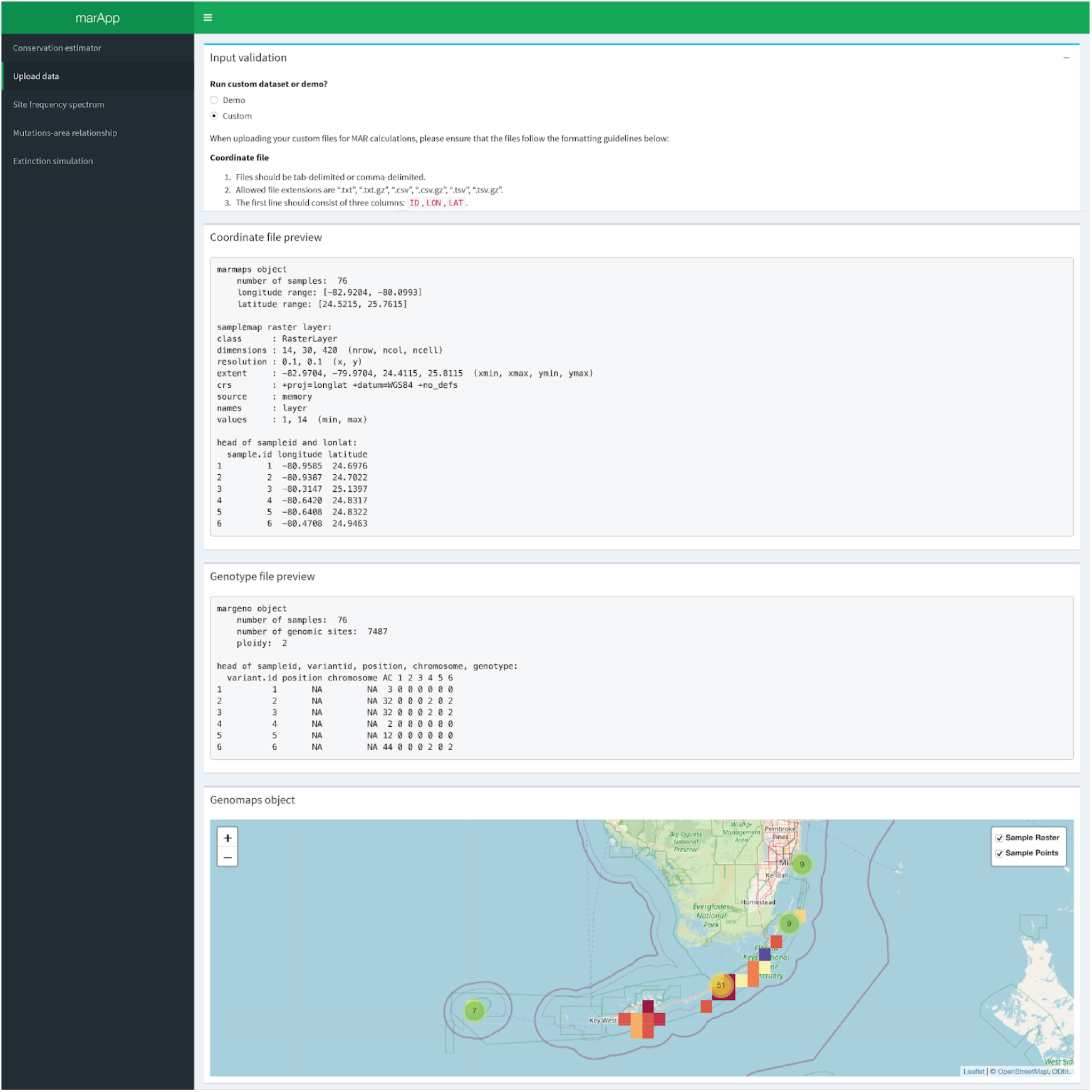
Visualization from the *marApp* “Upload data” tab using the Florida *Acropora* example dataset.

In the “Site Frequency Spectrum” tab, after selecting the species-abundance distribution (SAD) models to fit (defaults are log-normal and Fisher’s log-series distributions), select “Compute SFS and fit SAD”. The statistics for SAD model fits are provided, and the best-fit SAD model is automatically ranked on top. The log-likelihoods computed from SAD-predicted SFSs using the `ll_sfs` function are also provided. On the right, an interactive viewing of SFS computed from the genotype data (data), the population-genetics based neutral predictions (neutral), and selected SAD models are displayed. In this example, the log-series SAD (AIC = 51,365) fits the data better than the lognormal SADs (AIC = 54,709; **Fig. S4**).

In the “Mutations-area relationship” tab, the sampling schemes, genetic diversity metrics, area metrics and number of replicates can be customized and applied also to the “Extinction simulation” tab to ensure results comparability. After selecting “Calculate MAR/GDAR”, the summary table from the MAR sampling process is provided, together with the option to download outputs from the `MARsampling` function used to calculate the power-law relationship. To animate the sampling process, bounding boxes with various sizes used for MARsampling are overlaid with the sample map raster. Finally, the correlation of number of mutations and area sampled are interactively plotted, with the fitted *cA*^*z*^ curves overlaid as a gray line. Users can explore the power-law relationship on a log-scale and use different genetic diversity metrics. In this example, the scaling factor *z* for the number of segregating sites (*M*) and occupied grid cell area (*A*) is 0.33 (*p <* 0.001, *R*^*2*^ *=* 0.72; **Fig. S5)**.

In the “Extinction simulation” tab, after selecting “Simulate extinction”, the power-law relationship built from the extinction process is provided, together with the option to download outputs from the `MARextinction` function. To animate the simulated extinction process, at each extinction step, the raster cells that become unavailable to individuals are marked as black on the sample map raster. Towards the end of simulation, all cells become uninhabitable and the population is extinct. Finally, the percentage of habitable area lost is plotted with the percentage of genetic diversity remaining, with the MAR/GDAR-based theoretical predictions *(1 – a)*^*z*^ overlaid as a gray line. Users can explore the extinction prediction for different genetic diversity metrics. In this example, the predicted genetic diversity loss rate is *(1-a)*^*0*.*32*^ (**Fig. S6**).

The final step is the “Conservation estimator” tab, where users can use either the default scaling factor (*z =* 0.3), or the inferred scaling factor *z* for their species or regions of interest using a slider, to estimate the amount of genetic diversity loss or build habitat protection goals, and download the brief report generated. In this *Acropora* example, we sought to estimate the amount of genetic diversity loss using the scaling factor inferred from the “Extinction simulation” tab (*z =* 0.32) given that a combination of factors including disease, consistent ocean warming, and pollution has been threatening the survival of coral reef ecosystems (Cramer et al. 2020). In Florida Keys, healthy coral coverage has declined nearly 90% since the late 1970s (Gardner et al. 2003; Scott 2023), and *A. cervicornis* coverage reduced by 98% from 1983 to 2000 (Miller, Bourque, and Bohnsack 2002). Using the MAR relationship established above, we estimated that approximately 48% of *Acropora cervicornis* genetic diversity, measured in allelic richness, is still intact, and 52% of genetic diversity is already lost, assuming 90% short-term area loss (**Fig. 4**).

**Fig 4.**
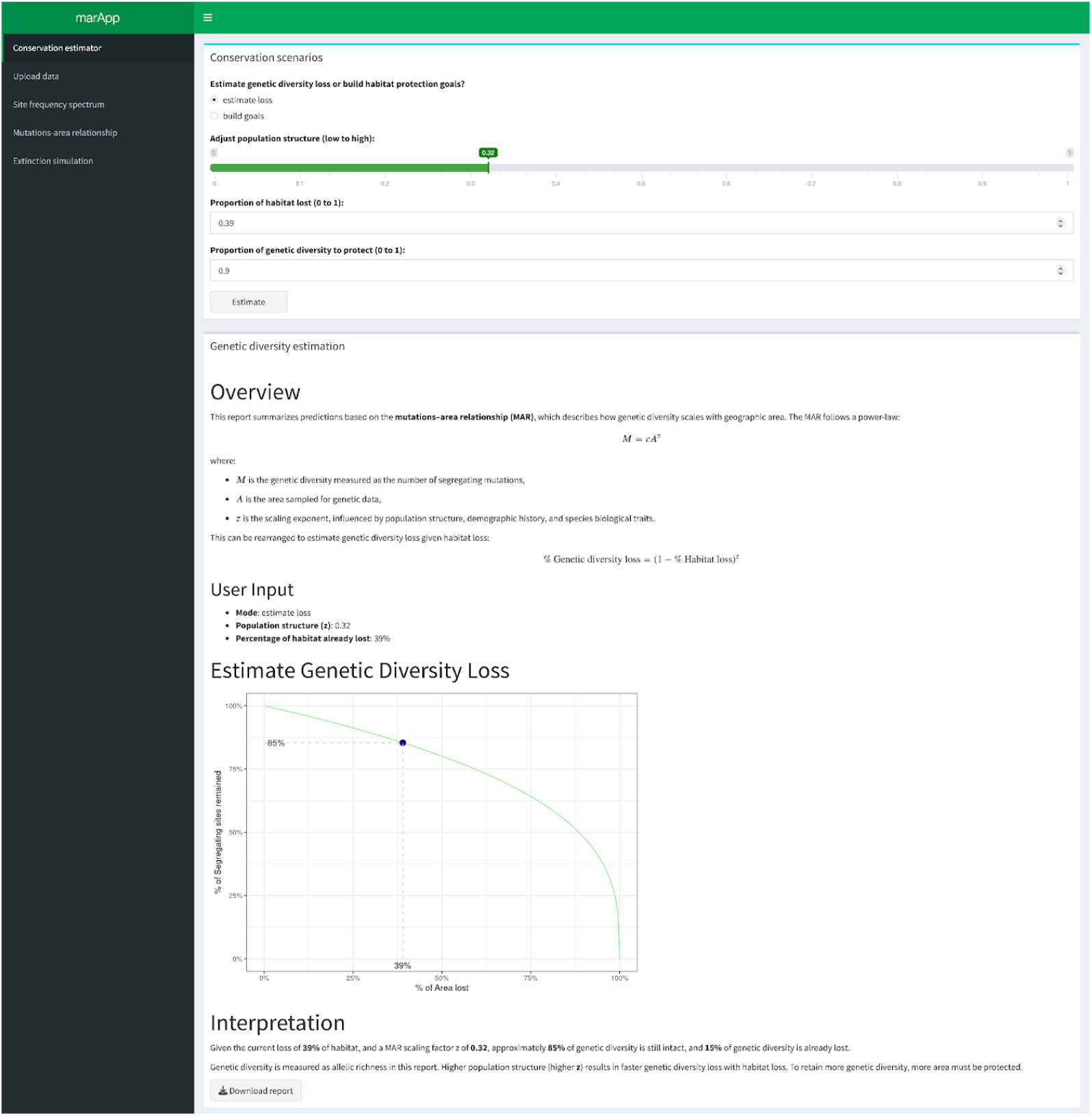
Screenshots from the *marApp* “Conservation estimation” tab using the Florida *Acropora* example dataset.

### Using the *marApp* to approximate genetic diversity loss without genetic data

The ultimate conservation advantage of the MAR framework is that it uses area to approximate genetic diversity loss in a simple percentage-based estimate. It may be thus helpful to understand the magnitude of genetic diversity loss for species that lack genetic data or for which this data is impossible or impractical to generate. In the *marApp*, the “Conservation estimation” tab can be utilized as a standalone tool for conservation planning. For example, in a recent multi-national evaluation of demographic-based genetic diversity indicators for 919 taxa, on average, 17.6% of wild populations have been lost per taxa, and 3% of taxa have lost 75% of their historical populations (Proportion of populations Maintained within species (PM) indicator in (Mastretta-Yanes et al. 2024)). Assuming population loss is proportional to habitat loss, under 75% area loss, genetic diversity, measured in allelic richness, reduces by 34% assuming a scaling factor of *z*_*MAR*_ ∼ 0.3 [34% = 1-(1-75%)^0.3^]. The scaling factor *z*_*MAR*_ describes the rate of genetic diversity loss, reflecting the spatial structure of genetic diversity. The scaling *z*_*MAR*_ approaches zero in a panmictic population with minimal spatial structure, and reaches one among completely isolated populations. Based on empirically inferred *z*_*MAR*_ values from 29 species (Exposito-Alonso et al. 2022; Mualim et al. 2024), *z*_*MAR*_ value of 0.3 likely captures large-scale trends of genetic diversity loss (Exposito-Alonso et al. 2022; Mualim et al. 2024). Using three representative *z*_*MAR*_ values, we offer interpretable genetic diversity loss estimates in nine countries with PM indicators (**Table 2**). For example, given the percentage population losses averaged across ∼100 species evaluated per country, if we assume a standard *z*_*MAR*_ of 0.3, the average remaining genetic diversity in Belgium is of 79% [79% = (45%)^0.3^], and at most 98% of genetic diversity is still maintained in countries including Japan, Mexico, and South Africa.

**Table 2.**
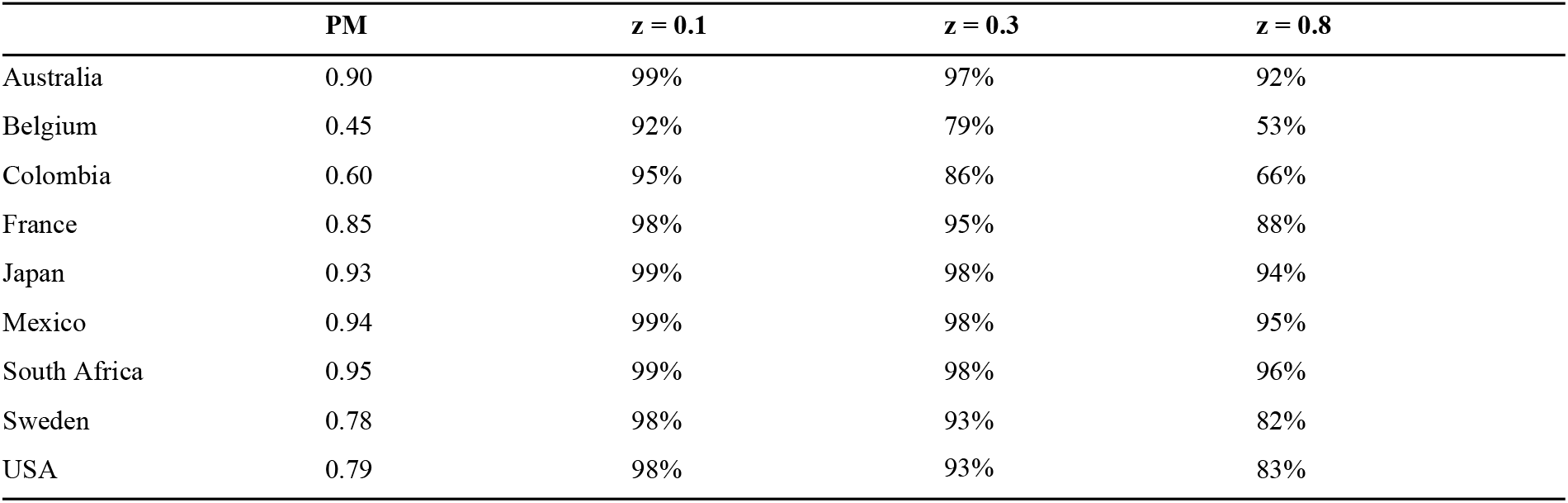
Estimated proportion of genetic diversity maintained within taxa for nine countries. Genetic diversity is measured in allelic richness. Estimates are based on short-term effects only, with more severe long-term effects expected.

| | PM | $z = 0.1$ | $z = 0.3$ | $z = 0.8$ |
| --- | --- | --- | --- | --- |
| Australia | 0.90 | 99% | 97% | 92% |
| Belgium | 0.45 | 92% | 79% | 53% |
| Colombia | 0.60 | 95% | 86% | 66% |
| France | 0.85 | 98% | 95% | 88% |
| Japan | 0.93 | 99% | 98% | 94% |
| Mexico | 0.94 | 99% | 98% | 95% |
| South Africa | 0.95 | 99% | 98% | 96% |
| Sweden | 0.78 | 98% | 93% | 82% |
| USA | 0.79 | 98% | 93% | 83% |

As the MAR/GDAR framework becomes adapted in more study systems with genetic data, we are only beginning to uncover the full extent of variation in *z* across species. To account for this uncertainty, we provide a reference table with different *z*_*MAR*_ and area-loss combinations (**Fig. S7**). Expanding collections of genetic data remains crucial, as it enables both the documentation of genetic diversity prior to potential loss and facilitates more robust estimations of *z*_*MAR*_ among species that are phylogenetically related or share similar life history traits (Leigh et al. 2021).

### Caution: long-term genetic diversity loss is larger than short-term *marApp* predictions

Importantly, in the “Conservation estimation” tab, we constrained the MAR framework to use only the number of segregating sites or allelic richness (*M*) as measures of genetic diversity, rather than nucleotide diversity (*θ*_*π*_), which is more commonly used in population genetics. This choice reflects that MAR/GDAR is a sampling formula describing contemporary genetic diversity, not a process-based temporal model capturing mutation-drift evolutionary dynamics over time (Mualim et al. 2024). Classically, average genetic diversity within a population has been well described by the mathematical formula: *θ*_*π*_ = 4 *N*_*e*_ *μ;* which captures the new mutations arising from mutation rate *μ* and the population size that can maintain them. During contemporary habitat loss, the census size (*N*_*c*_) declines instantly, while *θ*_*π*_ changes little because it still reflects pre-disturbance population size, *N*_*e*_. This creates a lag in *θ*_*π*_ responding to a population reduction, as shown in both the close-to-zero scaling factor *z*_*GDAR*_ ∼ 0.04 (Mualim et al. 2024). While allelic richness (*M*) responds quicker to recent population changes, it also suffers from this evolutionary lag (Mualim et al. 2024).

Due to this temporal lag of genetic diversity loss, we should not be feeling content on genetic diversity conservation. For example, we demonstrated above that in a scenario of 75% habitat loss, the allelic richness reduced less than the amount of habitat reduced by 34% (*z*_*MAR*_ = 0.3), and *θ*_*π*_ reduced even less by 15% (15% = 1 - (1-75%)^0.04^; *z*_*GDAR*_ *=* 0.04). However, in the long-term, when population reaches a new equilibrium, *θ*_*π*_ will eventually reflect the reduced effective population size, and reduce by 75% (z_GDAR_ ∼ 1 in the long-term (Mualim et al. 2024)) if habitats are not restored. This is akin to biodiversity conservation at the species richness level. While wildlife has already suffered much impact, only 1008 out of 169,420 (∼0.6%) species evaluated are classified as extinct or extinct in the wild (IUCN 2025). However, ecosystems are in a disequilibrium, 47,187 species are threatened by extinction, a phenomenon called “extinction debt”. Therefore, it is essential to protect as much genetic diversity as possible now and to act swiftly to reverse population declines, thereby preventing long-term genetic erosion and reducing the risk of future species extinction.

## Conclusion

The *mar* R package and *marApp* web portal fills an important gap in conservation, offering a theory-driven and percentage-based solution for integrating genetic diversity into conservation planning. By automating the analysis of mutations-area and other genetic diversity-area relationships (MAR/GDAR), *mar* makes it feasible to estimate genetic diversity loss under habitat reduction across species and landscapes. The user-friendly *marApp* further lowers the barrier for application by managers and policy makers, requiring no prior coding or genetics expertise. Through case studies in a global common species *Arabidopsis thaliana* and a local threatened species *Acropora cervicornis*, we show how MAR-based tools can generate quantitative forecasts, enabling alignment with targets in the Kunming-Montreal Global Biodiversity Framework. We caution that the *marApp* only provides estimates for genetic diversity in the short-term, and long-term genetic diversity loss is expected to be much more severe. While robust methods for long-term genetic diversity loss are under development for many species, it is essential to preserve as much genetic diversity as possible for the future. As the conservation field moves beyond species diversity toward preserving adaptive potential, *mar* and *marApp* provide an essential bridge between genomic data and actionable area-based decisions.

## Data availability

The *mar* package is available at https://github.com/meixilin/mar. The scripts used to build the *marApp* is available at https://github.com/meixilin/marApp. The data and scripts used to reproduce the *Arabidopsis* and *Acropora* examples are available at https://github.com/meixilin/marApp_paper. All genotype and coordinate data used in the examples are already publicly available.

## Acknowledgements

M.L. is supported by the David H. Smith Conservation Research Fellowship, and the Stanford Center for Computational, Evolutionary and Human Genomics. M.E.-A. is supported by the Office of the Director of the National Institutes of Health’s Early Investigator Award (1DP5OD029506-01), the U.S. Department of Energy, Office of Biological and Environmental Research (DE-SC0021286), by the U.S. National Science Foundation’s DBI Biology Integration Institute WALII (Water and Life Interface Institute, 2213983), by the Carnegie Institution for Science, the Howard Hughes Medical Institute, and the University of California Berkeley. The authors thank Alyssa Phillips and the MOILAB for helpful discussions.

## Author contributions

M.L. and M.E.-A. conceived the project. M.L. conducted research and wrote the first draft of the manuscript. M.E.-A. developed the beta version of *mar* package, K.M. performed package testing and O.S. provided scripts for analysis. All authors read, revised and approved the manuscript.

## Supplemental materials

### Supplemental tables

**Table S1.** Output from the mar step in `MARPIPELINE` for the *Arabidopsis* 1001G dataset. MAR were fitted using the *M = cA*^*z*^ function, from the `MARsampling(scheme = “random”)` output. “Mtype” denotes the four genetic diversity metrics considered: M: the number of segregating sites, E: the number of endemic segregating sites, thetaw: Watterson’s theta, and thetapi: nucleotide diversity. “Atype” denotes the two area calculation methods: A, the area of cells (km^2^) or Asq, the area of squares (º^2^). “c” and “z” are the inferred parameter values from *M = cA*^*z*^. “P(c)” and “P(z)” are the significant values for the inferred “c” and “z” values respectively. “R^2^” column shows the adjusted R^2^ value for each fitted genetic diversity and area relationship.

| Mtype | Atype | c | z | P(c) | P(z) | R<sup>2</sup> |
| --- | --- | --- | --- | --- | --- | --- |
| M | A | 7.25 | 0.50 | 7.84E-48 | 0.00E+00 | 0.97 |
| E | A | 5.62E-05 | 1.28 | 6.10E-07 | 0.00E+00 | 0.98 |
| thetaw | A | 3.04E-03 | 0.25 | 1.72E-36 | 3.55E-184 | 0.80 |
| thetapi | A | 1.43E-02 | 0.10 | 6.13E-49 | 2.41E-72 | 0.50 |
| M | Asq | 310.80 | 0.39 | 3.38E-08 | 5.11E-47 | 0.46 |
| E | Asq | 5.39 | 0.81 | 7.87E-02 | 7.28E-25 | 0.36 |
| thetaw | Asq | 2.52E-02 | 0.15 | 9.57E-37 | 8.24E-41 | 0.32 |
| thetapi | Asq | 3.29E-02 | 0.07 | 1.58E-95 | 2.08E-27 | 0.22 |

### Supplemental figures

**Fig S1.**
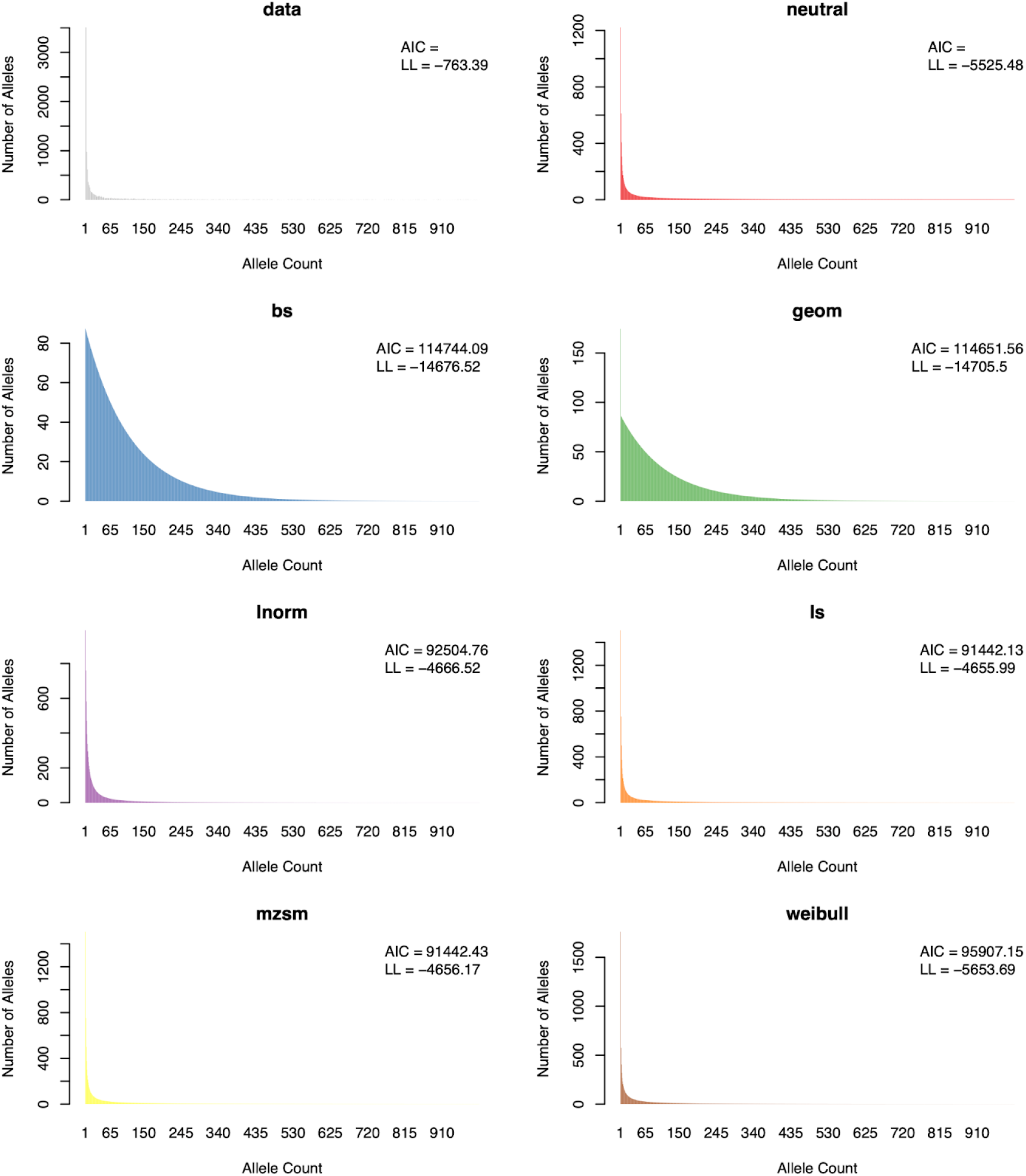
Output from `plot.sfs` in `MARPIPELINE` for the *Arabidopsis* 1001G dataset. The site frequency spectrum (SFS) of the data (gray, topleft), the expected SFS from neutral coalescence (red, topright), and the expected SFS from individual species abundance distributions (SAD). The AIC output is annotated for all SAD models, and the log-likelihood computed from the SFS is annotated for all SFSs.

**Fig S2.**
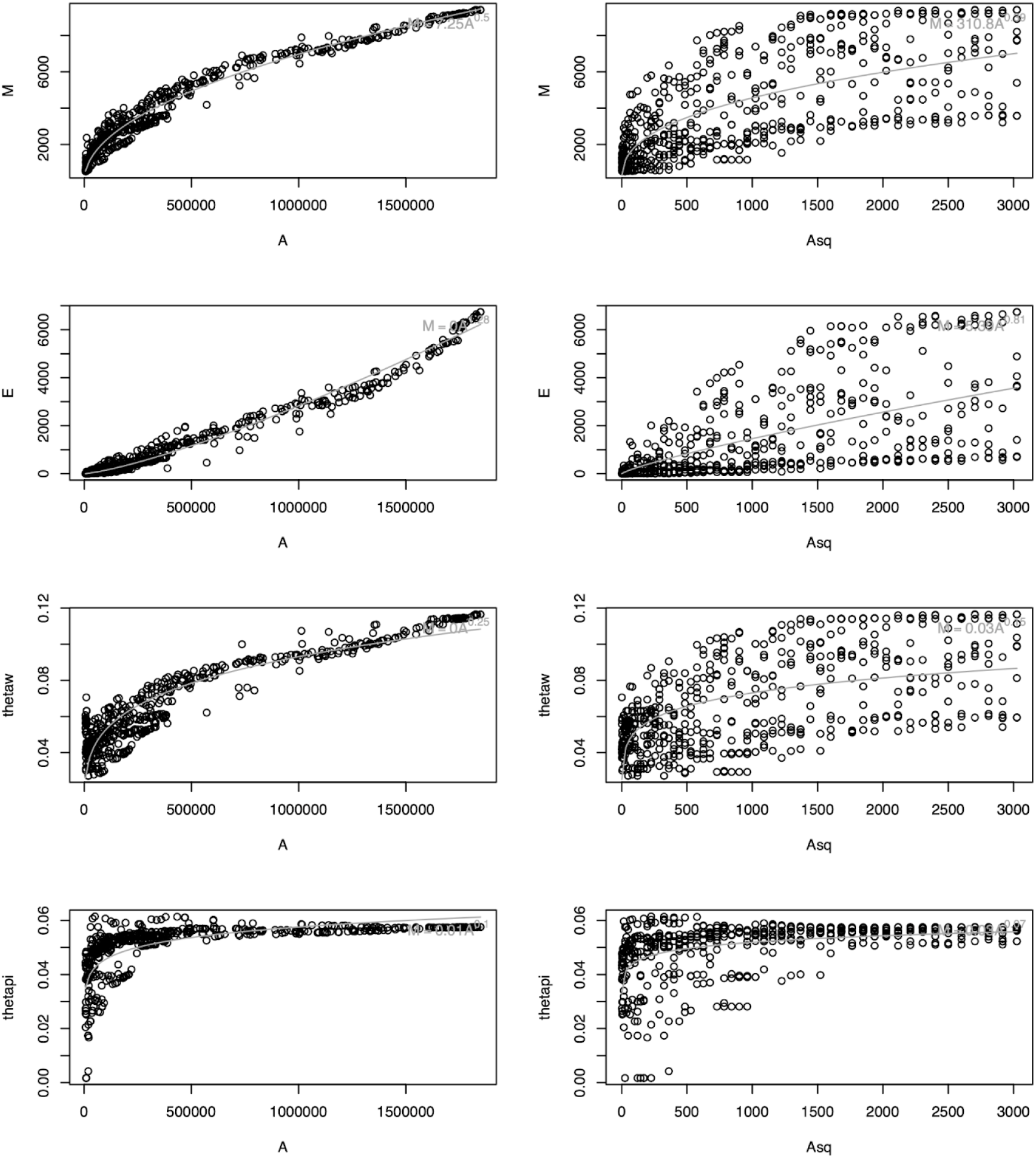
Output from `plot.marsamp` in `MARPIPELINE` for the *Arabidopsis* 1001G dataset. MAR were constructed from the `MARsampling(scheme = “random”)` output, using either *A*, the area of cells (km^2^, on the left), or *Asq*, the area of squares (º^2^, on the right). Four genetic diversity metrics were considered, from top to bottom, M: the number of segregating sites, E: the number of endemic segregating sites, thetaw: Watterson’s theta, and thetapi: nucleotide diversity. The fitted *M = cA*^*z*^ line is overlaid and the equation annotated on the top right in each panel.

**Fig S3.**
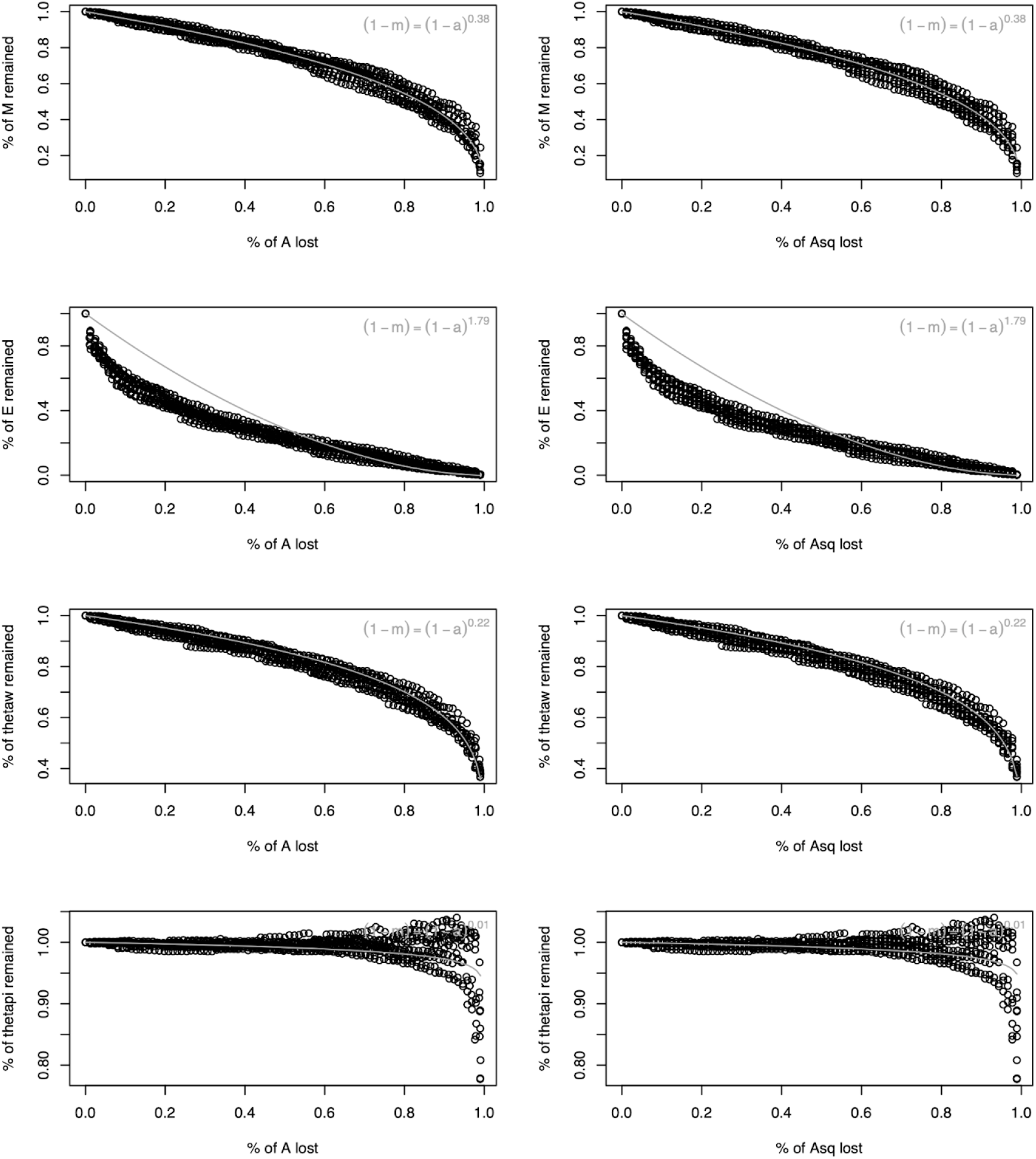
Output from `plot.marextinct` in `MARPIPELINE` for the *Arabidopsis* 1001G dataset.. Genetic diversity loss predictions were constructed from the `MARextinction(scheme = “random”)` output, using either *A*, the area of cells (km^2^, on the left), or *Asq*, the area of squares (º^2^, on the right). Four genetic diversity metrics were considered, from top to bottom, M: the number of segregating sites, E: the number of endemic segregating sites, thetaw: Watterson’s theta, and thetapi: nucleotide diversity. The fitted (*1 - m) = (1 - a)*^*z*^ line is overlaid and the equation annotated on the top right in each panel.

**Fig S4.**
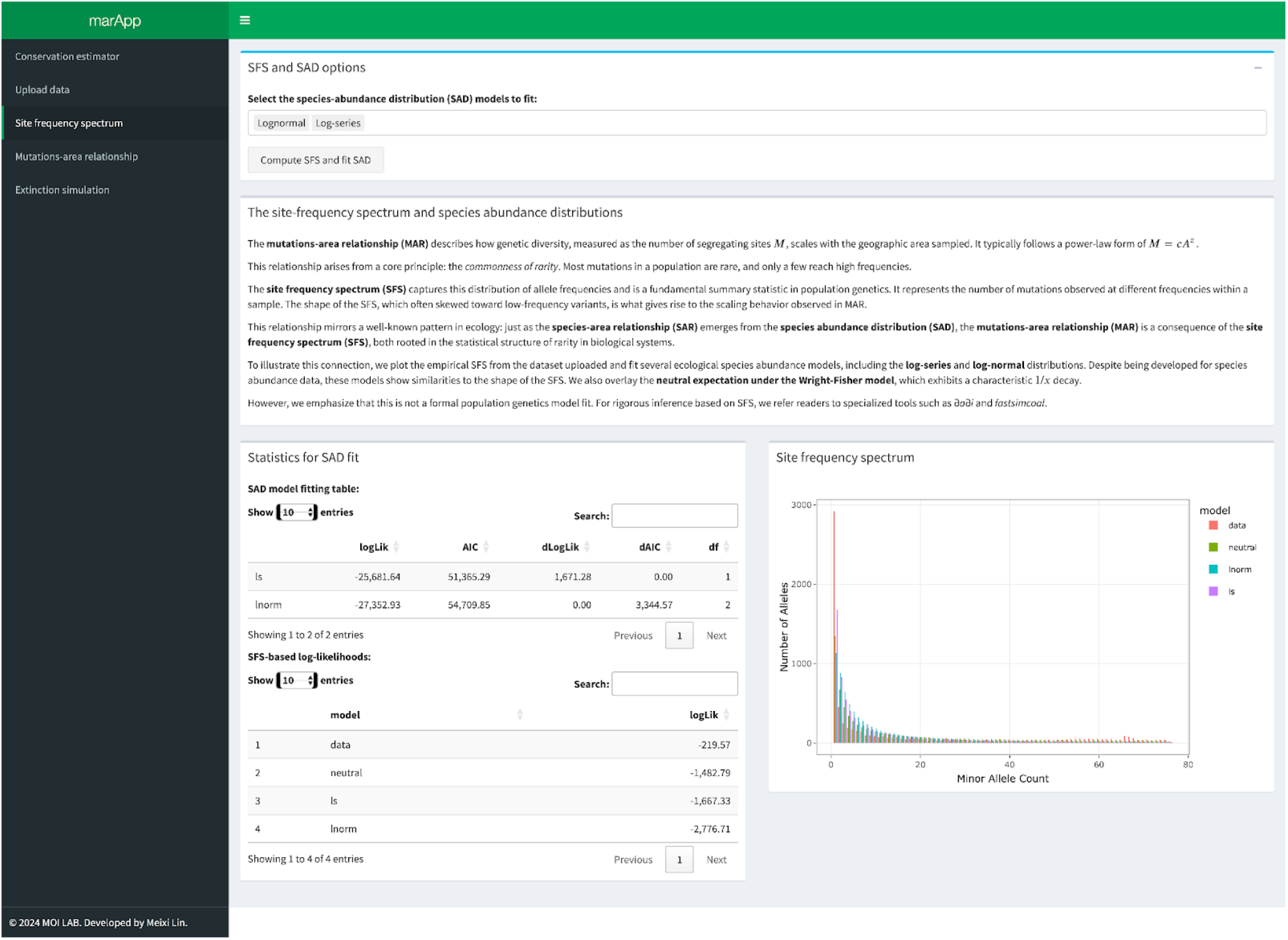
Screenshots from the *marApp* Site frequency spectrum tab using the Florida *Acropora* example dataset.

**Fig S5.**
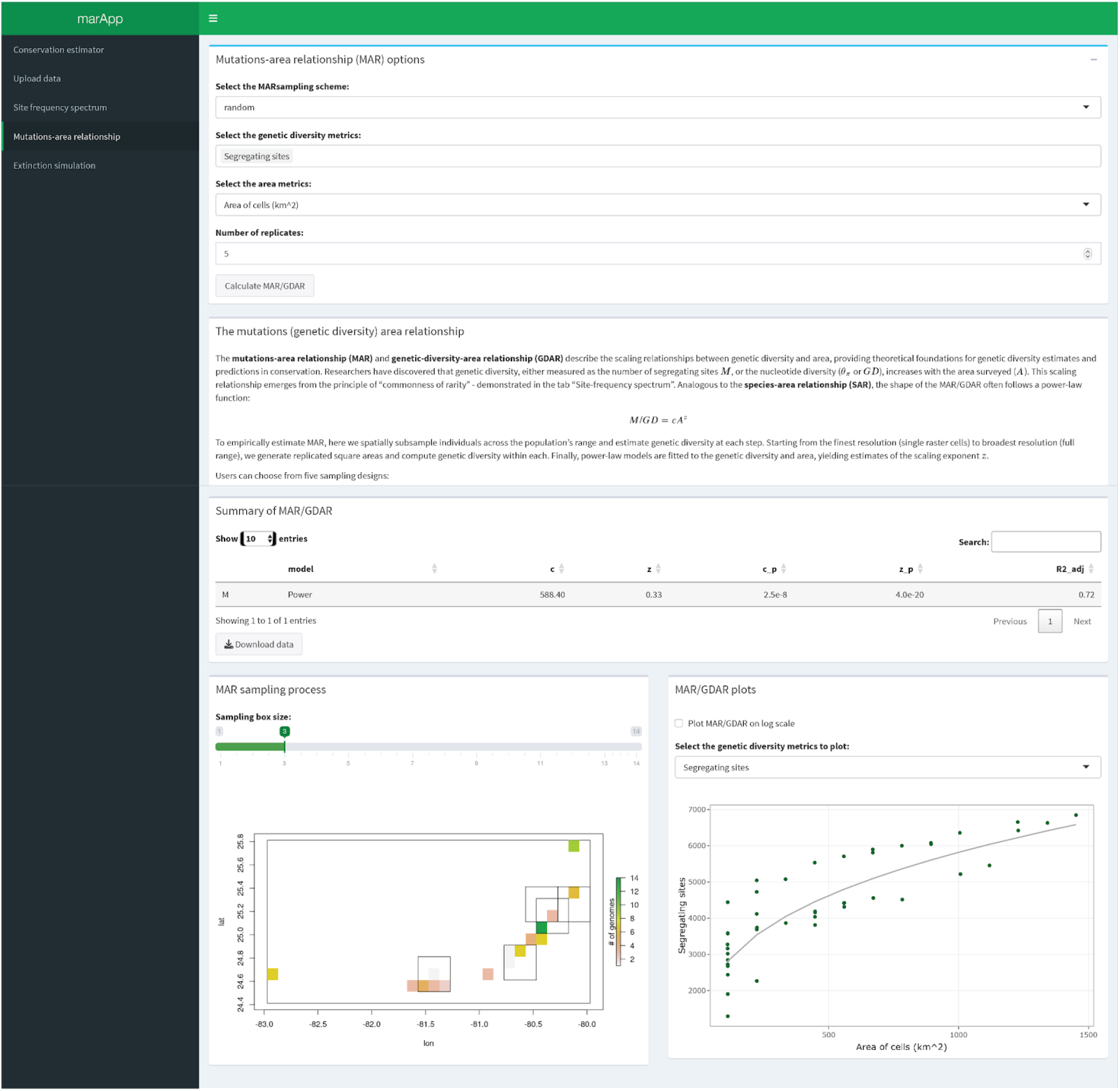
Truncated screenshots from the *marApp* Mutations-area relationship tab using the Florida *Acropora* example dataset.

**Fig S6.**
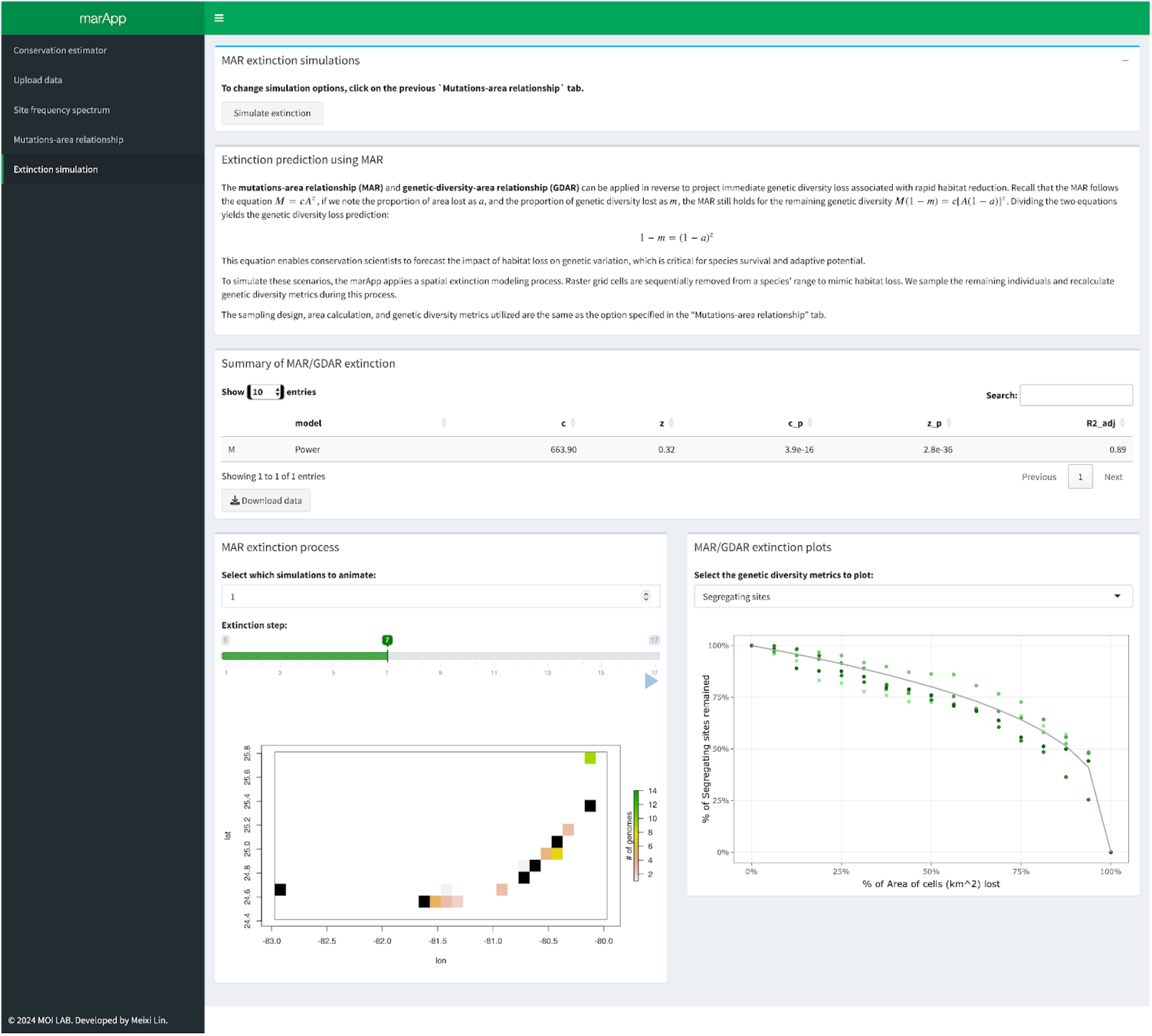
Truncated screenshots from the *marApp* Extinction simulation tab using the Florida *Acropora* example dataset.

**Fig S7.**
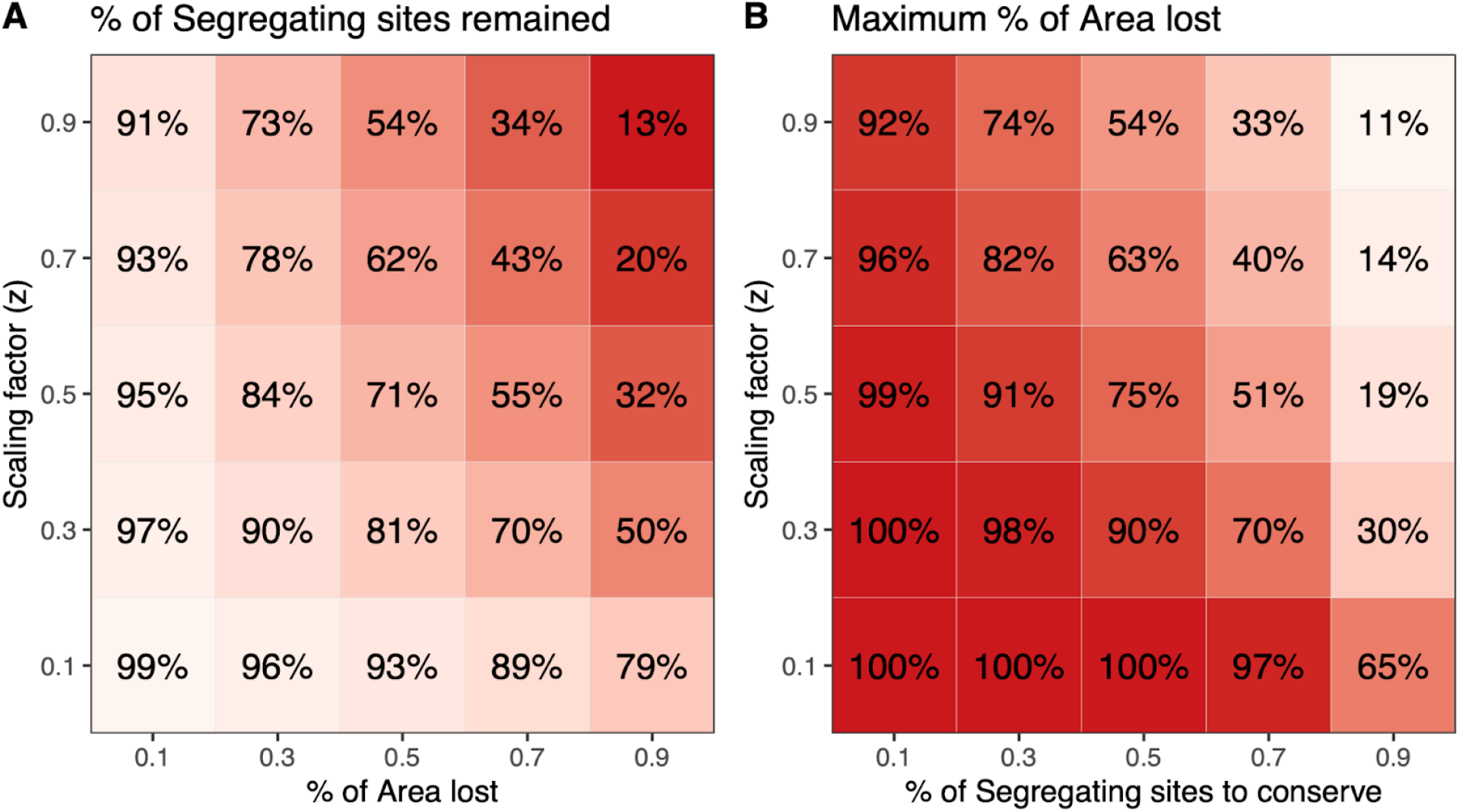
Reference table for approximating genetic diversity loss without genetic data. (A) Given a scaling factor (*z*), and a known habitat loss scenario, the approximate amount of genetic diversity remaining in the population is plotted, with darker color representing more genetic diversity loss. (B) Given a scaling factor (*z*), and a known genetic diversity conservation target, the maximum allowable percentage of area loss is plotted, with darker color representing more area loss. All estimates are based on the number of segregating sites (allelic richness) in a short-term scenario. Much more genetic diversity loss is expected in the long-term (Mualim et al. 2024).

## Notes

### Competing Interest Statement

The authors have declared no competing interest.

